# Item-specific delay activity demonstrates concurrent storage of multiple items in working memory

**DOI:** 10.1101/382879

**Authors:** David W. Sutterer, Joshua J. Foster, Kirsten C.S. Adam, Edward K. Vogel, Edward Awh

## Abstract

**Abstract:** A longstanding view holds that information is maintained in working memory (WM) via persistent neural activity that encodes the content of WM. Recent work, however, has challenged the view that *all* items stored in WM are actively maintained. Instead, “activity-silent” models propose that items can be maintained in WM without the need for persistent neural activity, raising the possibility that only a subset of items – perhaps just a single item – may be actively represented at a given time. While past studies have successfully decoded multiple items stored in WM, these studies cannot rule out an active switching account in which only a single item is actively represented at a time. Here, we directly tested whether multiple representations can be held concurrently in an active state. We tracked spatial representations in WM using alpha-band (8–12 Hz) activity, which encodes spatial positions held in WM. Human observers (male and female) remembered one or two positions over a short delay while we recorded EEG. Using a spatial encoding model, we reconstructed stimulus-specific working memory representations (channel tuning functions, CTFs) from the scalp distribution of alpha-band power. Consistent with past work, we found the selectivity of spatial CTFs was lower when two items were stored than when one item was stored. Critically, data-driven simulations revealed that the selectivity of spatial representations in the two-item condition could not be explained by models restricting storage to a single item at a time. Thus, our findings provide robust evidence for the concurrent storage of multiple items in visual working memory.

**Author Summary:** Working memory (WM) is a mental workspace where we temporarily hold information “online” in pursuit of our current goals. However, recent activity-silent models of WM have challenged the view that all items are held in an “online” state, instead proposing that only a subset of representations in WM – perhaps just one item – are represented by persistent activity at a time. To directly test a single-item model of persistent activity, we used a spatial encoding model to read out the strength of two representations from alpha-band (8–12 Hz) power in the human EEG signal. We provide direct evidence that both locations were maintained *concurrently*, ruling out the possibility that declines in stimulus-specific activity are due to storing one of two items in an activity-silent state.

## Introduction

Working memory (WM) is an “online” memory system that maintains information in a readily accessible state. A longstanding view is that WM representations are maintained via persistent, stimulus-specific delay activity. Visual features maintained in WM can be decoded from patterns of persistent neural activity in humans and nonhuman primates alike (Funahashi et al., 1989, 1993; Lara and Wallis, 2014; Harrison and Tong, 2009; Serences et al., 2009). In line with the known declines in behavioral performance as WM load increases (Luck and Vogel, 1997; Wilken and Ma, 2004; Zhang and Luck, 2008), the selectivity of stimulus-specific delay-period activity also declines with increasing memory load (Buschman et al., 2011; Emrich et al., 2013; Matsushima and Tanaka, 2014; Sprague et al., 2014, 2016). Thus, persistent activity has been thought to play a central role in WM maintenance (Sreenivasan et al., 2014).

Despite robust parallels to behavior findings, prior work does not firmly establish that multiple items are *concurrently* stored in an active state (i.e. represented by persistent neural activity). It has been postulated that reduced selectivity of neural representations with greater memory load occurs because competition between concurrently stored representations degrades the fidelity of those representations (Bays, 2014; Franconeri et al., 2013). However, recent “activity-silent” models of working memory have challenged the view that all items maintained in WM are supported by a persistent pattern of neural activity (e.g., continued firing of neurons), instead proposing that activity-silent memory mechanisms (e.g., rapid changes to synaptic weights) can support the short-term retention of information (Lewis-Peacock et al., 2012; Mongillo et al., 2008; Rose et al., 2016; Stokes, 2015; Wolff et al., 2017). In this view, when multiple items are stored in WM, they need not be concurrently represented by persistent activity. Instead, each item may transition between active and silent states, with only a single item in an active state at any given time. Indeed, recent work suggests that multiple items maintained with WM are activated serially (Bahramisharif et al., in press). Furthermore, other work suggests that when two locations must be attended, these locations are sampled sequentially (Fiebelkorn et al., 2013; Landau and Fries, 2012; vanRullen et al., 2007). Given past work positing a functional overlap between spatial WM and spatial attention (Awh and Jonides, 2001; Awh et al., 2006; Gazzaley and Nobre, 2012), sequential representation may also underpin the maintenance of multiple locations in spatial WM. Such a *switching model*, in which only one item is actively represented at once, also predicts an apparent decline in the fidelity of neural representations when mnemonic load is increased, because typical analyses aggregate data across multiple trials (e.g., Emrich et al., 2013; Sprague et al., 2014). Thus, if an item is represented for a smaller portion of the delay period when load increases, this could mimic the effects of competition between concurrently stored items.

Here, we provide a decisive test of whether persistent neural activity can concurrently represent multiple items. Human observers maintained the locations of one or two items over a brief delay period. We tested whether multiple locations are concurrently represented in oscillatory alpha-band (8–12 Hz) activity, which past work has shown precisely encodes spatial locations in WM (Foster et al., 2016, 2017). To this end, we used an inverted encoding model (lEM; Brouwer and Heeger, 2009; Sprague et al., 2015) to reconstruct representations of the remembered locations from the scalp distribution of EEG alpha-band power (Foster et al., 2016, 2017). Consistent with past work, we found that the selectivity of stimulus-specific activity was reduced when two items were remembered than when one item was remembered. Critically, however, when we simulated the expected selectivity of stimulus-specific activity during the two-item condition assuming a switching model (in which only one item was actively represented at once), we found that the observed selectivity in the two-item condition was reliably higher than predicted by a switching model. Thus, our findings provide robust evidence for the concurrent representation of at least two items in an active state.

## Results

### Behavior

Observers performed a spatial working memory task (Fig. 1A). On each trial, observers remembered the spatial position of one or two colored dots and reported the position of a cued item with a mouse click. We found that median response times (Fig. 1B) were slower for two-item trials (M = 1050 ms, *SD* = 245) than one-item trials (*M* = 1224, *SD* = 271), *t*(27) = 12.33, *p* < .0001. In line with past work (Luck and Vogel, 1997; Wilken and Ma, 2004; Zhang and Luck, 2008; Bays et al., 2009), memory performance declined as memory load increased from one to two items. We analyzed the recall data using a three-component mixture model (Bays et al., 2009) to estimate mnemonic precision (SD, higher values indicate lower precision), the probability that a stimulus was forgotten (*pGuess*), and the probability of reporting a non-target item (*pSwap*). We found that mnemonic precision was worse (i.e., *SD* was higher, Fig. 1C) when participants maintained two items (*M* = 6.83°, *SD* = 1.05) than when they maintained one item (*M* = 5.45°, *SD* = 1.01), *t*(27) = 15.66, *p* < .0001. We saw no reliable difference in the rate of guessing (Fig. 1D) between one-item (*M* = 0.12%, *SD* = 0.24) and two-item (*M* = 0.08%, *SD* = 0.16) trials, *t*(27) = -1.60, *p* = 0.121. Finally, observers’ rates of misreporting the location of the non-target item (Fig. 1E) on two-item trials (*M* = 0.23%, *SD* = 0.30) was reliably greater than zero, *t*(27) = 4.10, *p* = 0.0003. Note that the combined rate of guessing and swapping were very low (less than 1% for almost all observers). Thus, the primary change in behavior with memory load was the slowing of response times and reduction in precision of responses.

**Figure 1.**
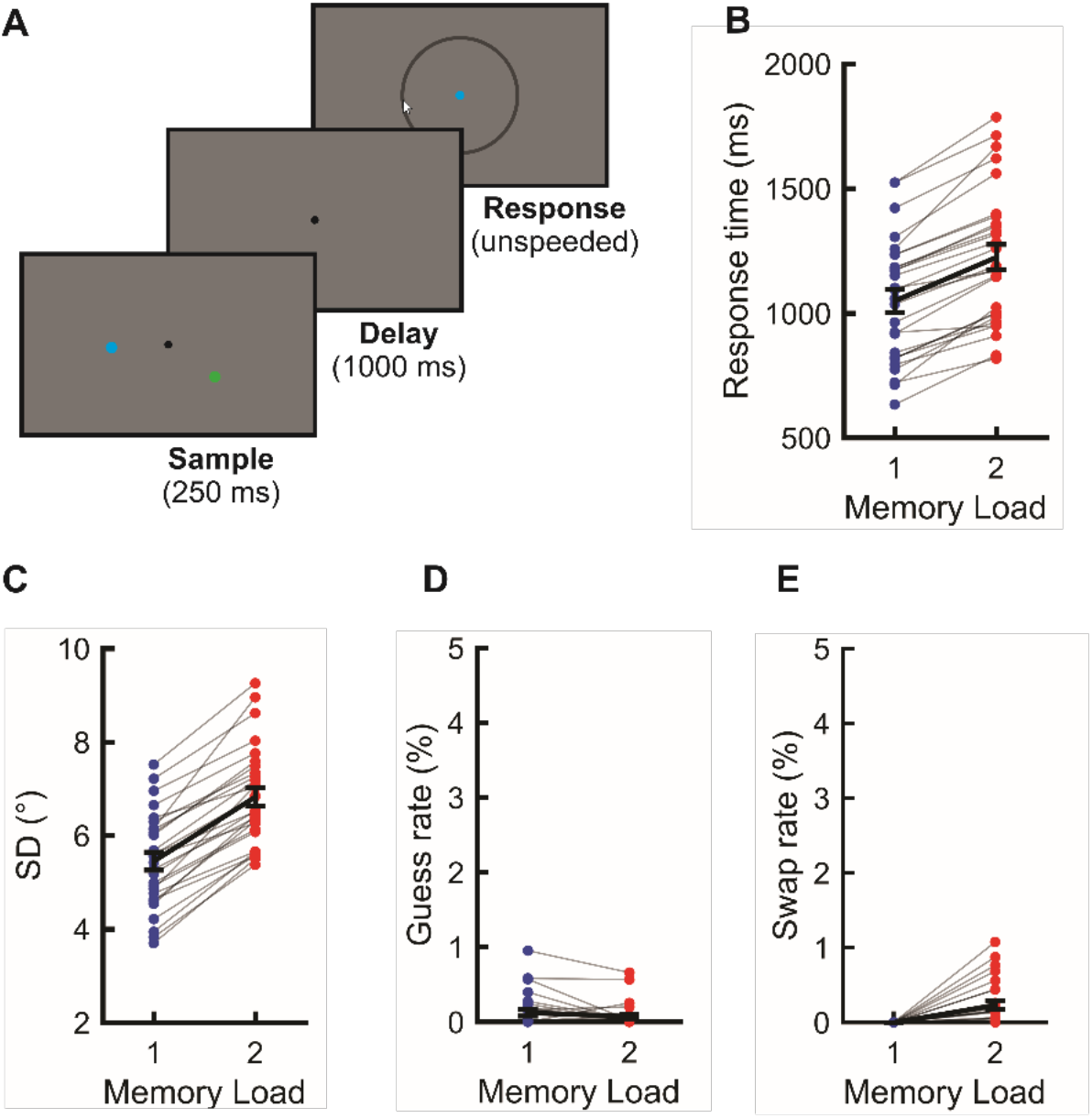
Experimental task and behavior. **A**, Observers saw a brief sample display (250 ms) that contained one or two colored dots. After a delay period (1Ũ00 ms), the fixation point changed color (blue or green) to indicate which item should be reported. Observers reported the angular position of the cued item as precisely as possible by mouse-click on the perimeter of a rim. **B**, Median response time (ms) as a function of memory set size (one vs. two items). Light grey lines represent individual observers. Black lines represent the mean. Error bars represent ±1 SEM. **C-E**, Parameter estimates obtained by fitting a three-component mixture model (Bays et al., 2009) to response errors as a function of memory set size. SD (**C**) reflects precision of responses (with higher values indicating worse precision), pGuess (**D**) estimates the probability that the observer produced a random response (i.e., a guess), and pSwap (**E**) estimates the probability that an uncued item was m¡sreported instead of the cued-item. Note that pSwap is necessarily zero for the single-item condition.

### Alpha-band representations of space degrade with increased memory load

To test how online representations change with increased memory load, we examined oscillatory alpha-band (8–12 Hz) activity, which encodes spatial representations that are maintained in WM (Foster et al., 2016, 2017). We used an inverted encoding model (IEM; Brouwer and Heeger, 2009, 2011; Sprague et al., 2015) to reconstruct spatial representations encoded by alpha-band activity (Foster et al., 2016). Our encoding model assumes that alpha-band power measured at each scalp electrode reflects the activity of a number of spatially tuned channels (or neuronal populations), each tuned for a different position in the visual field (Fig. 2A and 2B). In a training phase, we estimated the relative contributions of the spatial channels to each electrode on the scalp (called the “channel weights”) using a subset of trials during the spatial WM task (Fig. 2C). Then, in a test phase, using an independent subset of trials, we used these channel weights to estimate the responses of the spatial channels given the pattern of alpha-band power across the scalp. The resulting profile of responses across the spatial channels (called channel-tuning functions, or CTFs) reflects the spatial selectivity of alpha-band activity measured by EEG. We performed this analysis at each time point throughout the trial, which allowed us to test whether active spatial representations were maintained throughout the delay period.

**Figure 2.**
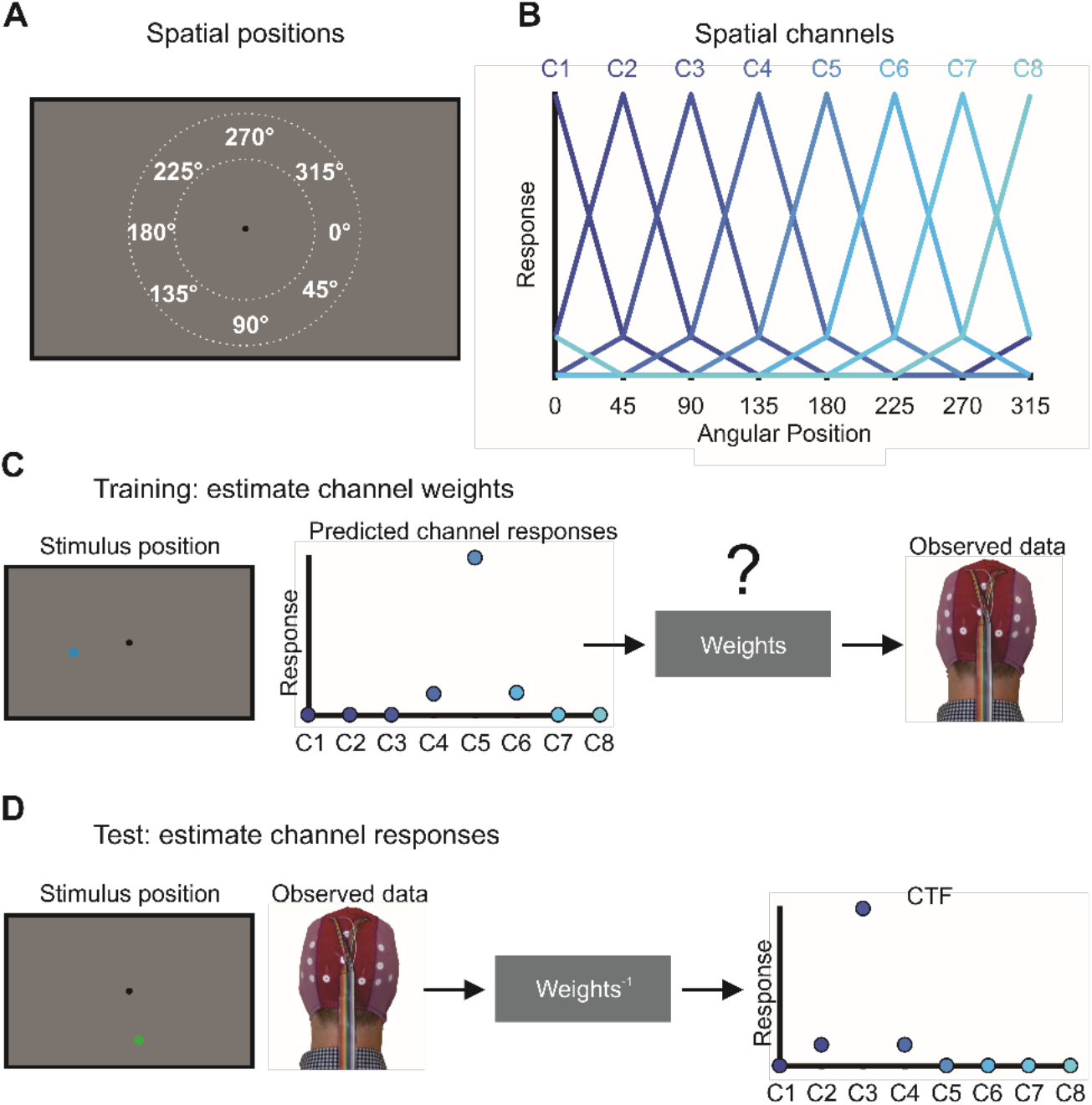
Inverted Encoding model procedure for reconstruction spatial channel tuning function (CTFs) ***A***, In our experiment, the sample stimuli could appear along an isoeccentric band of positions around the fixation point. We categorized stimuli as belonging to one of eight location bins centered at 0°, 45°, 90°, etc. Each position bin spanned a 45° wedge of positions (e.g., 22.5° to 67.5° for the bin centered at 45°). ***B***, we modeled oscillatory power at each electrode as the weighted sum of eight spatially selective channels (C1-C8), each tuned for the center of one of the eight position bins shown in (A). Each curve shows the predicted response of one of the channels across the eight position bins (i.e., the “basis function”). ***C***, In the training phase, we used the predicted channel responses, determined by the basis functions shown in (B), to estimate a set of channel weights that specified the contribution of each spatial channel to the response measured at each electrode. The example shown here is for a stimulus occupying the location bin centered at 180°. ***D***, In the test phase, using an independent set of data, we used the channel weights obtained in the training phase to estimate the profile of channel responses given the pattern of activity across the scalp. The resulting CTF reflects the spatial selectivity of population-level oscillatory activity, as measured with EEG. The example shown here is for a stimulus occupying the location bin centered at 90°. For details, see Materials and Methods.

**Figure 3.**
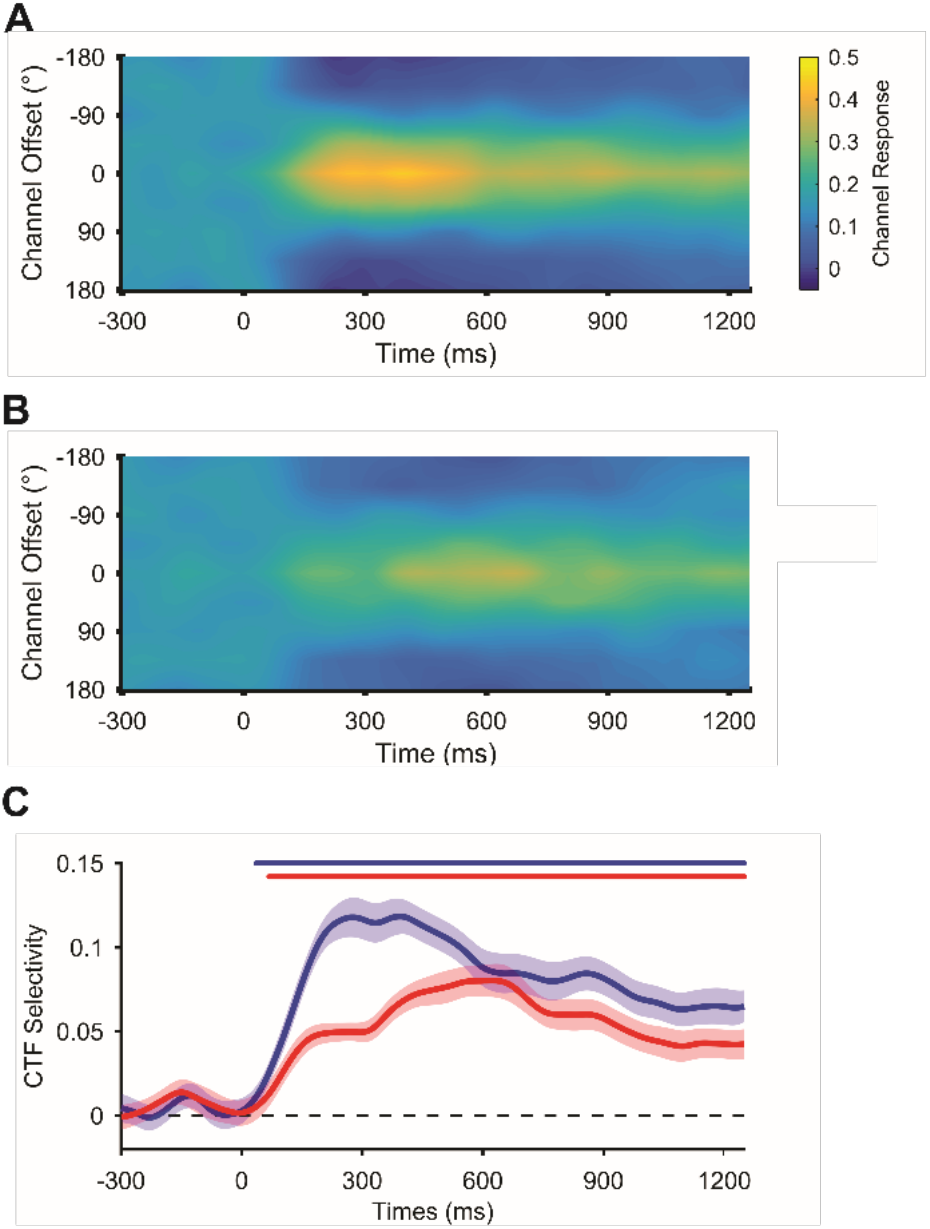
Spatial alpha-band CTFs as a function of memory load. Average alpha-band CTF in the one-and two-item conditions (***A*** and ***B***, respectively). ***C***, The spatial selectivity of alpha-band CTFs across time (measured as CTF slope, see Materials and Methods) as a function of memory load. The blue (one-item) and red (two-item) markers at the top of the panel indicate the period of above-chance selectivity obtained using a cluster-based test. CTF selectivity was reliably higher in the one-item condition than in the two-item condition. The shaded error bars reflect ±1 bootstrapped SEM across observers.

To examine how WM load affects alpha-band representations of the remembered positions, we reconstructed CTFs for the both the one- and two-item conditions (see Materials and Methods). We observed a clear spatially selective CTF, with a peak response in the channel tuned for the remembered location (a channel offset of 0°), which persisted throughout the delay period in both the one-item and two-item conditions (Fig. 3A and 3B). Figure 3C shows the spatial selectivity of the CTFs seen in each condition across time (measured as CTF slope, see Materials and Methods). Cluster-based permutation tests revealed that spatial selectivity of alpha-band CTFs was reliably above zero throughout the delay period for both the one- and two-item conditions (*p* < .05, corrected for multiple comparisons; see markers at the top of Fig. 3C). Next, we compared CTF selectivity for the one-item and two-item conditions. A resampling test confirmed that delay-period CTF selectivity (averaged from 250 to 1250 ms after stimulus onset) was reliably lower (*p* < .0001) for two-item trials (*M* = .059, *SD* = .033) than for one-item trials (*M* = .087, *SD* = .046). Importantly, we also found that this difference was reliable (*p* = .004) during a late window (800–1250 ms) when alpha-band representations are unlikely to be affected by stimulus-driven activity. Thus, as memory load increases, there is a decline in spatially-selective alpha-band activity that tracks the stored locations.

### Alpha-band activity concurrently encodes two spatial representations

Consistent with past work (Buschman et al., 2011; Emrich et al., 2013; Sprague et al., 2014, 2016), we observed that spatially specific alpha-band activity deteriorates as memory load increases. However, declines in stimulus-specific activity with increasing load can be explained in two ways. On the one hand, the observed decrease in selectivity might reflect the loss of memory fidelity due to competition between representations when multiple stimuli are maintained in an active state (Bays, 2014; Franconeri et al., 2013). On the other hand, activity-silent models of WM (Stokes, 2015), highlight the possibility that only a single item is maintained in an active state at any given moment, while the other items stored in WM are represented in an activity-silent state. Under this account, the apparent persistent activity in the two-item condition (Fig. 3B) can be explained if the two items in WM switch in and out of an active state. (Fig. 4A). Critically, this switching account asserts that with a memory load of two items, each item can only be represented a maximum of 50% of the time, on average. To test whether this switching account can explain the CTFs seen during the two-item condition, we simulated the CTF selectivity expected under a switching account. To this end, we generated CTFs from the single-item condition but randomized the position labels for 50% of the trials (see Materials and Methods). We then compared CTF selectivity seen during the two-item condition with the CTF selectivity expected under the switching account. We reasoned that if the switching account was correct, we should see no difference between CTF selectivity for two-item trials and for the simulated switching conditions. However, if we observed a higher CTF selectivity for the observed two-item data, we could conclude that alpha activity reflects the simultaneous maintenance of multiple locations in WM during a given trial.

**Figure 4.**
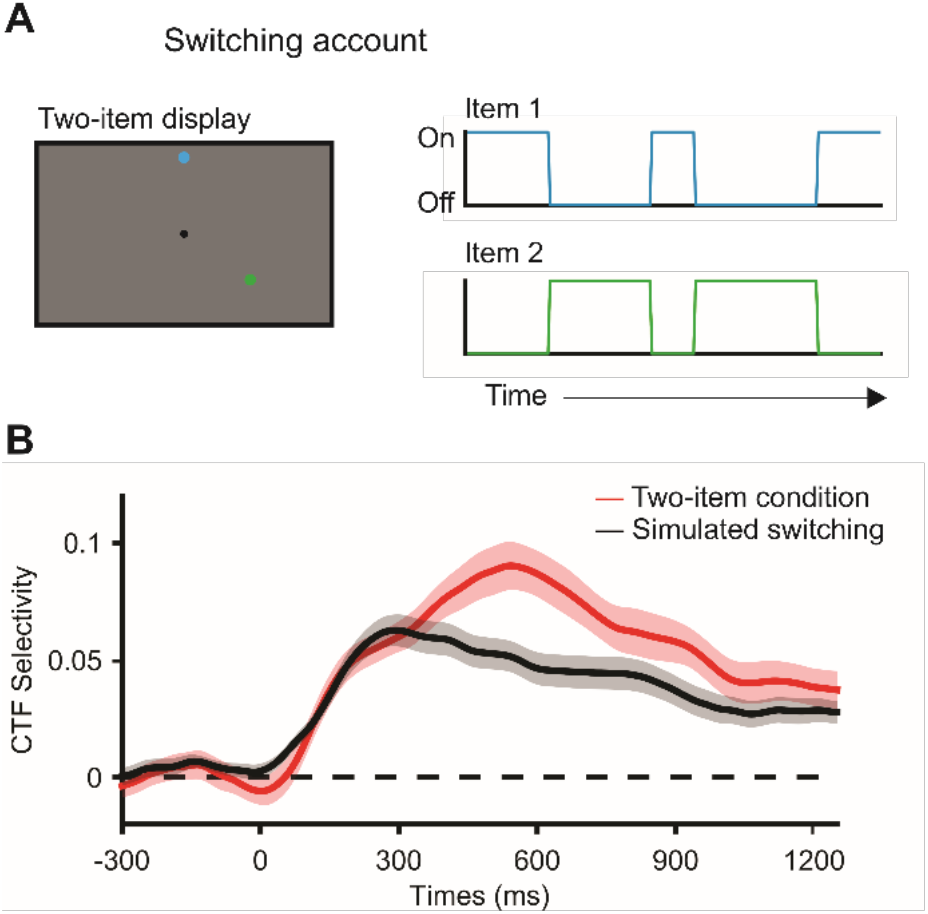
Alpha-band activity concurrently represents two spatial positions. ***A***, If only one item can be represented by alpha-band activity at a time, then in the two-item condition the items might alternate between an active and an activity-silent state such that only one item is actively represented at once. We used data from the single-item condition to simulate the expected CTF based on this switching account. This account holds that each item is represented 50% of the time (on average). Thus, we simulated switching between items by randomizing the position labels for 50% of trials, ***B***, Spatial selectivity of alpha-band CTFs across time (measured as CTF slope) for the two-item condition (red) and for simulated switching (black). CTF selectivity was reliably higher during the two-item condition (red) than for simulated switching (black). The shaded error bars reflect ±1 bootstrapped SEM across observers.

Figure 4B shows CTF selectivity across time for the two-item condition and for simulated switching based on the one-item data. We found that CTF selectivity was higher throughout the delay period (averaged from 250 to 1250 ms after stimulus onset) for the two-item condition (*M* = .063, *SD* = .036) than expected based on the switching account (*M* = .043, *SD* = .030). A resampling test revealed that this difference was reliable (*p* < .0001). We also observed a reliable difference when we restricted our analysis to a window late in the delay period (800-1250 ms) to minimize the contribution of stimulus-driven activity (*p* = .001). This analysis provides definitive evidence that multiple locations are simultaneously represented by alpha-band activity, and that observed decreases in CTF selectivity as memory load increases reflect a decline in the quality of concurrently stored representations rather than rapid switching between active representations.

### The frequency of oscillations that encode spatial representations does not change with memory load

The decline in the spatial selectivity of alpha-band activity with increasing memory load suggests that the fidelity of the spatial representations decreased as memory load increased. However, another possibility is that the remembered locations were represented by a different frequency band when memory load increased. To test this possibility, we performed the lEM analysis separately for the one-item and two-item conditions (i.e., both training and testing within each condition; see Materials and Methods) across a range of frequencies (4–50 Hz). We conducted a cluster-corrected permutation test to identify reliable clusters of above-chance CTF selectivity. Consistent with our past work (Foster et al., 2016), we observed a burst of spatially specific activity across a range of frequencies (4–25 Hz).

However, only alpha-band activity (8–12 Hz) tracked the remembered position(s) throughout the delay period (Fig. 5A and 5B). An overlay plot of spatially specific frequencies in both one-item and two-item trials revealed a strikingly similar frequency profile later during the delay period, when stimulus-driven activity has subsided (Fig. 5C). These findings show that the frequency of oscillatory activity that encodes spatial representations does not change with memory load. Thus, the decrease observed in the spatial selectivity of alpha-band CTFs with increasing memory load reflects a decline in spatially selective activity rather than a shift in the frequency of spatially selective oscillations.

**Figure 5.**
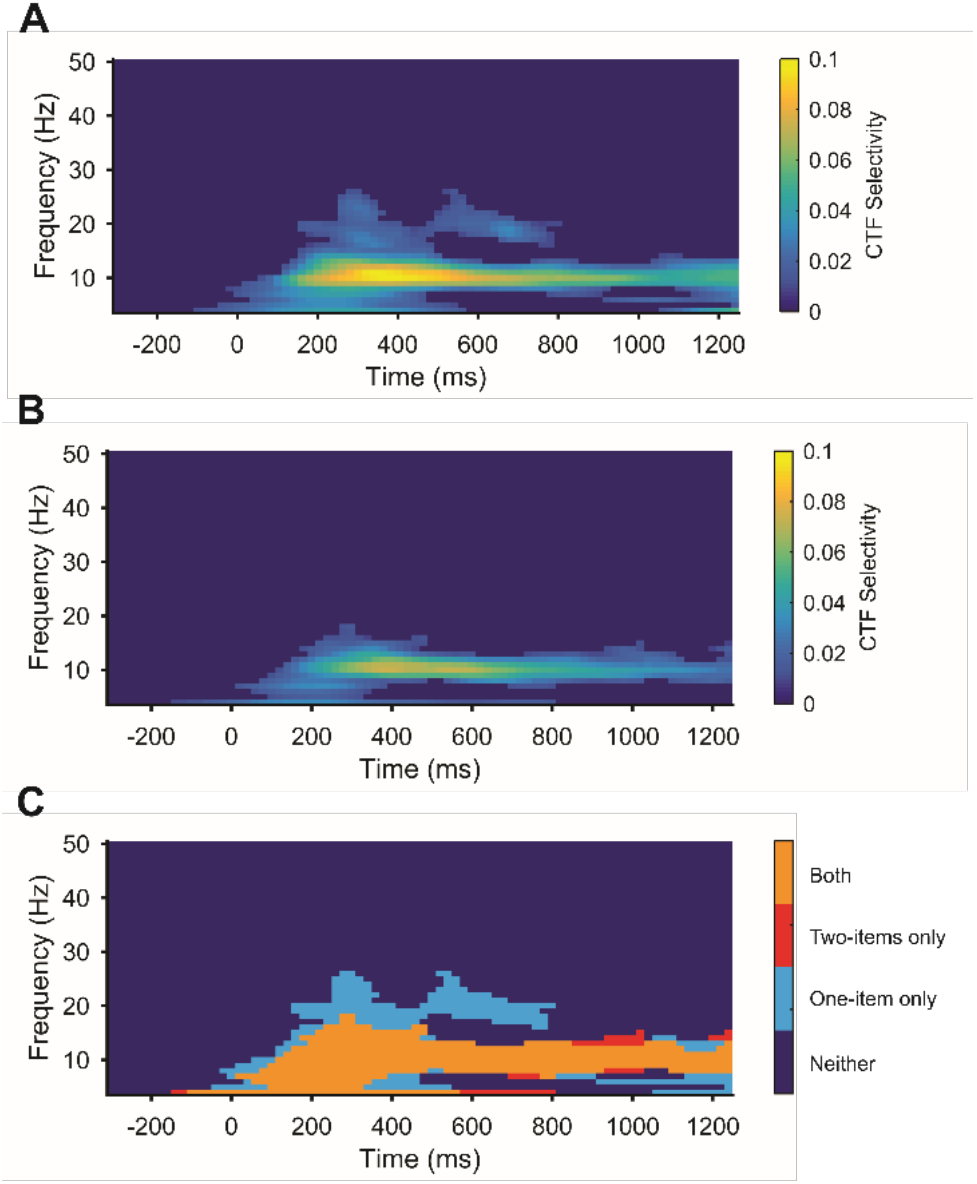
***A-B***, Selectivity of spatial CTFs (measured as CTF slope) reconstructed from the scalp distribution of oscillatory power across a broad range of frequencies for the one-item (***A***) and two-item (***B***) conditions. Points with no reliable CTF selectivity as determined by the cluster-corrected permutation test are set to dark blue. ***C***, Overlay plot marking the clusters of reliable selectivity in the one-item condition (light blue), two-item condition (red), and both (orange).

### Ruling out chunking of locations when two items are stored in WM

In our two item condition, observers precisely reported each of the two locations and we saw clear spatially specific delay activity in this condition. However, past work has suggested that multiple items may be stored in WM by maintaining a single “chunk” in WM rather than independent representations of each item (e.g., Huang and Awh, 2018). We reasoned that if observers maintained the two locations as a single chunk, then the pattern of spatially selective alpha-band activity on the scalp should not generalize from the one-item condition to the two-item condition. In contrast, if observers maintained two independent spatial locations, we expected that pattern of alpha-band activity corresponding to each location should generalize from the one-item condition to the two-item condition. To test between these two possibilities, we trained the IEM on one-item trials, and tested on two-item trials. This analysis revealed robust reconstructions of the remembered locations (Fig. 6), providing clear evidence that the spatially selective pattern of alpha-band activity generalizes from the one-item condition to the two-item condition. This finding supports the view that observers maintained two distinct spatial representations in spatial working memory.

**Figure 6.**
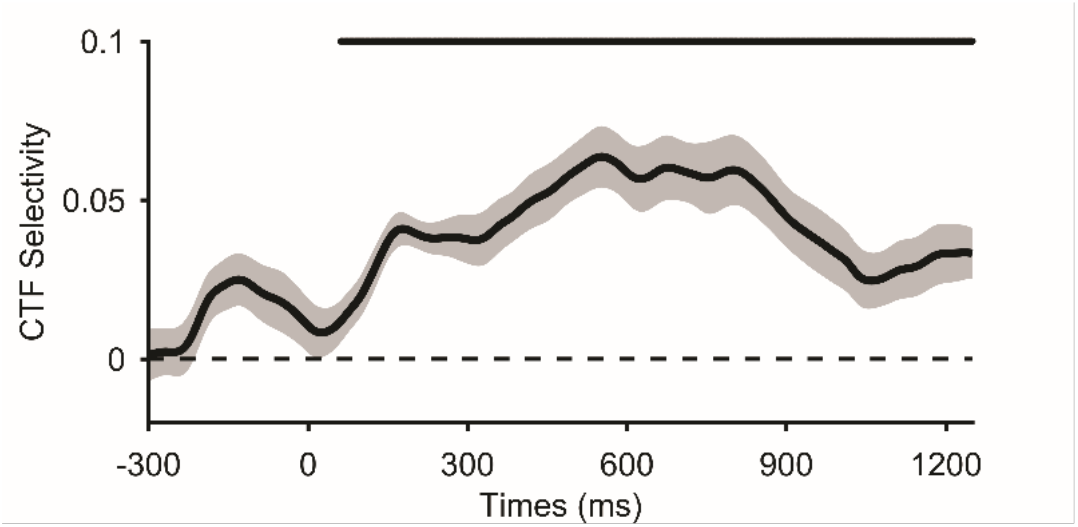
The spatial selectivity of alpha-band CTFs across time (measured as CTF slope, see Materials and Methods) in the two-item condition when the IEM was estimated using the one-item condition. The marker at the top of the panel indicate the period of above-chance selectivity obtained using a cluster-based test. The shaded error bars reflect ±1 bootstrapped SEM across observers.

## Discussion

A longstanding view is that the maintenance of information in WM is realized via persistent neural activity that encodes the content of WM. In support of this view, past work has shown that stimulus-specific patterns of delay-period activity track visual features maintained in WM (Funahashi et al., 1989, 1993; Foster et al., 2016; Harrison and Tong, 2009; LaRocque et al, 2017; Serences et al., 2009; for review, see Sreenivasan et al., 2014). However, recent work has challenged stimulus-specific activity as the sole mechanism that supports maintenance in WM, instead proposing that other activity-silent mechanisms can also support the retention of information (Lewis-Peacock et al., 2012; Stokes, 2015; Rose et al., 2016; Wolff et al., 2017). This activity-silent account warrants a re-evaluation of the evidence that all items maintained in WM are represented in stimulus-specific patterns of activity. The prevailing view has been that multiple items are concurrently represented by stimulus-specific delay activity. Indeed, past work finds that increased memory load results in a decline in the stimulus-specific patterns of activity during WM maintenance (Buschman et al., 2011; Emrich et al., 2013; Sprague et al., 2014, 2016), mirroring behavioral declines in performance. However, the activity-silent account raises the possibility that only a single item is actively represented at a time, and that the *apparent* representation of multiple items is seen because typical analyses aggregate data across trials, averaging together periods when an item is represented and when it is not represented.

Here, we tested such a switching account, in which only one item was actively represented at once. We used an inverted encoding model (IEM) combined with EEG measurements of oscillatory alpha-band (8–12 Hz) power to track remembered spatial locations during a WM task in which human observers remembered one or two spatial locations. Consistent with previous fMRI and non-human primate unit recordings that found that stimulus-specific activity declines with WM load (e.g., Buschman et al., 2011; Emrich et al., 2013; Sprague et al., 2014), we found that the spatial selectivity of alpha-band activity declined as memory load increased. We tested the switching account by simulating selectivity expected under a switching model in which only one item was actively represented at a time. We found that the spatial selectivity of alpha-band CTFs during two-item trials was greater than predicted by models that restrict active storage to only one item at a time, even when rapid switching is allowed. Thus, our findings provide clear evidence that two items can be represented concurrently in an active state.

In principle, our switching simulation can also be applied to existing MRI studies that have examined stimulus-specific activity as a function of memory load (e.g., Emrich et al., 2013; LaRocque et al., 2017; Sprague et al., 2014, 2016). However, the BOLD (blood oxygen level dependent) signal measures the hemodynamic response as a proxy for neural activity rather than measuring neural activity directly. Thus, evidence for concurrent representations in the BOLD signal does not necessarily indicate concurrent representations in neural activity, especially given the slow time course of the hemodynamic response. For example, if two items were switched in and out of an active state (e.g., on the order of hundreds of milliseconds), this would likely produce concurrent representations of both items in the BOLD signal. In contrast, we directly measured alpha-band activity. Thus, we can be confident that both items are concurrently represented by neural activity.

The encoding model approach – which allowed us to directly assay stimulus-specific activity – was critical to test whether two items were concurrently represented in an active state. Past work has identified load-dependent neural signals that track the number of items maintained in WM, reaching an asymptote around WM capacity. For example, Vogel and Machizawa (2004) reported a lateralized EEG signal (called contralateral delay activity or CDA) that tracked the number of items held in WM, reaching an asymptote around 3-4 items. Similarly, Todd and Marois (2004) found that delay activity in the intraparietal sulcus showed a similar load-dependent profile. We and others have viewed these load-dependents signals as tracking the number of active representations in WM (Luck and Vogel, 2013; Luria et al., 2016). However, neural activity that scales with the number of items in memory array does not necessitate that multiple active representations are maintained concurrently in WM. For example, under an activity-silent account in which only one item is actively represented at once, load-dependent activity could track the rate at which items are switched in and out of an active state. When more items must be maintained, items may be switched in and out of an active state more rapidly, which might produce activity that tracks memory load. In contrast, because we directly examined stimulus-specific activity, we were able to provide unambiguous evidence that multiple representations were actively represented at once. It may well be that load-dependent signals track the number of active representations. However, more work is needed to test this claim. If load-dependent signals track the number of active representations in WM, then load-dependent activity should predict the number of active representations in WM.

At first glance, our finding that two spatial representations can be concurrently represented in an active state seems inconsistent with spatial attention studies that have provided evidence for rhythmic sampling when multiple items or locations must be attended or stored (e.g., Bahramisharif et al., in press; Busch and Van Rullen, 2010). However, the switching account that we tested is a stringent version of rhythmic sampling in which items alternate between “on” and “off” states. Our results do not rule out all classes of rhythmic sampling models, but they do constrain these models. Specifically, our data demonstrate that even if there are orderly rhythms that describe when a given item is best represented, a complete model must allow for the concurrent representation of multiple items. Thus, if the primary active representation switches from one item to the other, then this handoff must be done in such a way that both items are actively represented during the switch.

Finally, evidence that two active representations can be maintained concurrently is relevant to a long-standing debate about the nature of representations in WM. Embedded process models characterize memory as a common storage space with different levels of activation corresponding to short-term memory and long-term memory (for review, see LaRocque et al., 2014). One key distinction between competing embedded process models is the number of items that can be represented in an active state in WM. The proposed number of items ranges from a strict limit of one item (McElree, 2006; Oberauer, 2002) to 3-4 items (Cowan, 1995; Luck and Vogel, 2013), and competing models are each able to account for many aspects of behavioral performance (LaRocque et al., 2014). Our findings provide clear evidence that two items can be actively represented concurrently, disconfirming the class of models that posit a strict one-item limit on the number of items that can be actively represented.

## Materials and Methods

### Participants

Forty-one volunteers participated in the experiment for monetary compensation ($15/hr). Participants were between 18 and 35 years old, reported normal color vision and normal or corrected-to-normal visual acuity, and provided informed consent according to procedures approved by the University of Chicago Institutional Review Board. The sample included both male and female participants. We excluded participants from the final sample if fewer than 450 trials per condition remained after discarding trials contaminated by recording or ocular artifacts (see Artifact Rejection). Eight observers were excluded because too few trials remained after artifact rejection. Data collection was terminated early for four observers because the data were unusable due to excessive artifacts, and for one observer because of a fire alarm. The final sample included 28 observers with an average of 581 (*SD* = 75) trials for one-item trials and 596 (*SD* = 69) trials for two-item trials.

### Apparatus and Stimuli

We tested participants in a dimly lit, electrically shielded chamber. Stimuli were generated using Matlab (MathWorks, Natick, MA) and the Psychophysics Toolbox (Brainard, 1997; Pelli, 1997), and were presented on a 24” LCD monitor (refresh rate: 120 Hz, resolution: 1080 × 1920 pixels) at a viewing distance of ~100 cm. Stimuli were rendered against a gray background.

### Task procedure

Participants performed a spatial delayed-estimation task in which they remembered the spatial position of one or two sample stimuli (see Fig. 1A). Participants initiated each trial with a spacebar press. Each trial began with a fixation point (0.2° in diameter) presented for 500-800 ms. Next, a memory array which comprised one or two sample stimuli was presented for 250 ms. Each stimulus was a blue or green circle (0.2° in diameter, equated for luminance) centered 4° of visual angle from the fixation point. On one-item trials, the sample stimulus was blue or green. On two-item trials, one stimulus was blue and the other was green. Participants were instructed to remember the spatial position of the sample stimuli as precisely as possible. The angular position of each stimulus around the fixation point was sampled from eight position bins, each spanning a 45° wedge of angular positions (bins were centered at 0°, 45°, 90°, and so forth, see Fig. 2A), with jitter added to cover all 360° of possible locations to prevent categorical coding of stimulus location. On two-item trials, the position bins that each stimulus occupied were fully counterbalanced across trials for each observer. Thus, the position bin that one stimulus occupied was random with respect to the other, which allowed us to reconstruct spatial CTFs for each stimulus independently. When both stimuli occupied the same position bin, their exact position within the bin was constrained so that the two items were separated by at least 0.2° of visual angle. The memory array was followed by a 1000-ms delay period during which only the fixation point remained on screen. Finally, after the delay period, a cursor appeared at fixation and the fixation point turned blue or green to indicate which stimulus should be reported. Participants report the remembered location of the probed stimulus by using a mouse to click on the perimeter of a probe ring (8° in diameter, 0.2° thick). The color of the probed item (green or blue) was pseudo-randomized across trials and conditions such that each color was probed on 50% of trials for each condition. Before starting the task, participants completed a brief set of practice trials to ensure that they understood the task.

### Electrophysiology

We recorded EEG activity from 30 active Ag/AgCl electrodes mounted in an elastic cap (Brain Products actiCHamp, Munich, Germany). We recorded from International 10-20 sites: FP1, FP2, F7, F3, Fz, F4, F8, FT9, FC5, FC1, FC2, FC6, FT10, T7, C3, Cz, C4, T8, CP5, CP1, CP2, CP6, P7, P3, Pz, P4, P8, O1, Oz, O2. Two additional electrodes were affixed with stickers to the left and right mastoids, and a ground electrode was placed in the elastic cap at position FPz. Data were referenced online to the right mastoid and re-referenced offline to the algebraic average of the left and right mastoids. Eye movements and blinks were also monitored using electrooculogram (EOG), recorded with passive Ag/AgCl electrodes. Horizontal EOG was recorded from a bipolar pair of electrodes placed ~1 cm from the external canthus of each eye. Vertical eOg was recorded from a bipolar pair of electrodes placed above and below the right eye. Data were filtered online (low cut-off = .01 Hz, high cut-off = 80 Hz, slope from low- to high-cutoff = 12 dB/octave), and were digitized at 500 Hz using BrainVision Recorder (Brain Products, Munich, German) running on a PC. Impedance values were kept below 10 kΩ.

### Eye Tracking

We monitored gaze position using a desk-mounted EyeLink 1000 Plus infrared eye-tracking camera (SR Research, Ontario, Canada). Gaze position was sampled at 500 Hz, and data were obtained in remote mode (without a chin rest). We obtained usable eye-tracking data for 19 out of 28 participants.

### Artifact Rejection

We visually inspected the segmented EEG data for artifacts (amplifier saturation, excessive muscle noise, and skin potentials), and inspected EOG for ocular artifacts (blinks and eye movements). For observers with usable eye tracking data, we also inspected the gaze data for ocular artifacts. We discarded trials contaminated by artifacts. We discarded electrode FT9 and FT10 for all observers because we obtained poor quality data (excessive high-frequency noise) at these sites for most observers. Data from one or two electrodes were discarded for four participants because of excessive high-frequency noise or sudden steps in voltage which occurs when an electrode is damaged. The discarded electrodes for each participant were: T7; F3; CP6 and C4; and P8. For the analysis of gaze position, we further excluded trials in which the eye tracker was unable to detect the pupil, operationalized as any trial in which the horizontal or vertical gaze position was more than 15° from fixation. At most 15 trials per observer were rejected due to this reason. For most observers (15 of 19), no trials were excluded for this reason.

Removal of trials with ocular artifacts was effective. Variation in grand-averaged HEOG as a function of the remembered stimulus position was < 3.6 μV for both one-item trials and < 1.7 μV for two-item trials. Given that eye movements of about 1° of visual angle produce a deflection in the HEOG of ~16 μV (Lins et al., 1993), the residual variation in the HEOG corresponds to variations in eye position of < 0.23°. Analysis of the subset of participants (19 participants) for whom we obtained usable gaze position data corroborates the HEOG data obtained from all participants. Variation in grandaverage horizontal gaze position as a function of remembered stimulus position was < 0.12° for one-item trials and < 0.07° for two-item trials. For comparison, for these participants the variation in the average HEOG as a function of remembered stimulus position was < 3.1 μV for one-item trials and < 1.6 μV for two-item trials.

### Time-frequency Analysis

To calculate frequency specific activity at each electrode we first band-pass filtered the baselined raw EEG data using EEGLAB (“eegfilt.m”, Delorme and Makeig, 2004). For alpha-band analyses, the data were band-pass filtered between 8 to 12 Hz. For our exploratory analysis of a broad range of frequencies, we band-pass filtered the data in 1-Hz bands from 4 to 50 Hz (i.e., 4-5 Hz, 5-6 Hz, etc.). We applied a Hilbert transform (MATLAB Signal Processing Toolbox) to the band-pass-filtered data to obtain the complex analytic signal. Instantaneous power was calculated by squaring the complex magnitude of the complex analytic signal. To reduce computation time for the IEM analysis across time and frequency, we down-sampled the matrix of power values to one sample every 20 ms. We down-sampled power values (i.e., after filtering and applying the Hilbert transform) so that down-sampling did not affect the calculation of power.

### Inverted Encoding Model

Following our prior work (e.g., Foster et al., 2016, 2017), we used an inverted encoding model (IEM; Brouwer and Heeger, 2009, 2011; Sprague and Serences, 2013; for review, see Sprague et al., 2015) to reconstruct spatially selective channel-tuning functions (CTFs) from the topographic distribution of oscillatory power across electrodes. We assumed that the power at each electrode reflects the weighted sum of eight spatially selective channels (i.e., neuronal populations), each tuned for a different angular location (Fig. 2A). We modeled the response profile of each spatial channel across angular locations as a half sinusoid raised to the twenty-fifth power:

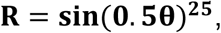

where θ is angular location (0–359°), and *R* is the response of the spatial channel in arbitrary units. This response profile was circularly shifted for each channel such that the peak response of each spatial channel was centered over one of the eight positions corresponding to the eight positions bins (0°, 45°, 90°, etc., see Fig. 2B).

An IEM routine was applied to each time point in the alpha-band analyses and to each time-frequency point in the time-frequency analyses. We partitioned our data into independent sets of training data and test data (see the Training and Test Data section). This routine proceeded in two stages (training and test). In the training stage (Fig 2C), training data (*B_1_*) were used to estimate weights that approximate the relative contribution of the eight spatial channels to the observed response measured at each electrode. Let *B_1_* (*m* electrodes × *n_1_* measurements) be the power at each electrode for each measurement in the training set, *C_1_* (*k* channels × *n_1_* measurements) be the predicted response of each spatial channel (determined by the basis functions, see Fig. 2B) for each measurement, and *W* (*m* electrodes × *k* channels) be a weight matrix that characterizes a linear mapping from “channel space” to “electrode space”. The relationship between *B_1_, C_1_*, and *W* can be described by a general linear model of the form:

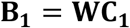

The weight matrix was obtained via least-squares estimation as follows:

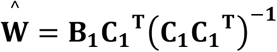

In the test stage (Fig. 2D), we inverted the model to transform the observed test data *B_2_* (*m* electrodes × *n_2_* measurements) into estimated channel responses, *C_2_* (*k* channels × Λ *n_2_* measurements), using the estimated weight matrix, 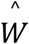, that we obtained in the training phase:

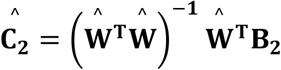

Each estimated channel response function was then circularly shifted to a common center, so the center channel was the channel tuned for the position of the probed stimulus (i.e., 0° on the “Channel Offset” axes). We then averaged these shifted channel-response functions across the eight position bins to obtain a CTF.

Finally, because the exact contributions of each spatial channel to each electrode (i.e., the channel weights, *W*) likely vary across participants, we applied the IEM routine separately for each participant, and statistical analyses were performed on the reconstructed CTFs. This approach allowed us to disregard differences in the how location-selective activity is mapped to scalp-distributed patterns of power across participants, and instead focus on the profile of activity in the common stimulus or “information” space (Sprague et al., 2015).

### Training and Test Data

For the IEM procedure, we partitioned artifact-free trials into independent sets of training data and test data for each observer. Across all analyses, we partitioned the trials into three independent sets. When portioning the trials into these sets, we equated the number of trials for each location in each set. Because of this constraint, some excess trials were not assigned to any set. Thus, we used an iterative approach to make use of all available trials. For each iteration, we randomly partitioned the trials into sets (as just described), and performed the IEM procedure on the resulting training and test data. Therefore, the trials that were not included in any block were different for each iteration. We averaged the resulting channel-response profiles across iterations. This iterative approach reduced noise in the resulting CTFs by minimizing the influence of idiosyncrasies that were specific to any given assignment of trials to blocks. For analyses focused on alpha-band power, we performed 50 iterations. For analyses across a wide range of frequencies (which is a time-consuming procedure), we performed 10 iterations.

Once trials were assigned to the three sets, we averaged across trials for each stimulus location bin to obtain a matrix of power values across all electrodes for each location bin (electrodes × locations, for each time point). We used a leave-one-out cross-validation routine such that two of these sets served as the training data, and the remaining matrix served as the test data. We applied the IEM routine using each of the three matrices as the test data, and the remaining two as the training set. The resulting CTFs were averaged across the three test sets. Different analyses require that the data for one and two item trials are partitioned into training and test sets differently depending on the goal of the analysis. In the following sub-sections, we outline how data were partitioned into training and test sets for each analysis.

### Comparing CTF selectivity across conditions

In two analyses, we tested whether CTF selectivity varied across condition. In the first analysis, we tested whether CTF selectivity differed as a function of memory load (see Fig. 3). In the second analysis, we compared CTF selectivity between the two-item condition and a condition that simulated switching using the one-item condition (see Fig. 4). These two analyses were assigned to training and test sets the same way. The only difference between the analyses was that for the simulated switching analysis, we randomized the position labels for half of the one-item trials after assigning trials to training and test sets. When comparing CTF properties across conditions, it is important to estimate a single encoding model that is then used to reconstruct CTFs for each condition separately. If this condition is not met, then it is difficult to interpret differences in CTF selectivity between conditions because these might result from differences between the training sets (i.e., how the model is estimated; for further discussion of this issue, see Sprague et al., 2018). We achieved this by estimating the encoding model using a training set consisting of equal trials from each condition. Specifically, we partitioned data for each condition into three sets (as described above, with the additional constraint that the number of trials per location in each set was also equated across conditions). We obtained condition-neutral training data by combining data across the two conditions before averaging, resulting in two training sets that included equal numbers of trials from each condition. We then tested the model of the remaining set of data for each condition separately. Thus, we used the same training data to estimate a single encoding model, and varied only the test data that was used to reconstruct CTFs for each condition.

### Time x Frequency analysis

In another analysis, we tested whether the range of frequencies that carried location-specific information varied as a function of memory load (see Fig. 5). For this analysis, we trained and tested the IEM within each condition separately. We trained and tested within each condition separately because we were interested in whether the frequencies that track the remembered location(s) differed between the conditions, a mixed training set would not be optimal to detect which frequencies do carry location-specific information in either condition. Thus, we partitioned data from each condition into three sets. As we did for the other analyses, we then trained the model using two out of three sets and tested the model using the one remaining set. Each of the three sets for each condition was held out as the test set. Critically, because the IEM was trained and tested within condition, we estimated a separate encoding model for each condition. Thus, this analysis maximizes our sensitivity to differences in the frequencies carrying spatial information between conditions; however, it is difficult to interpret any difference in CTF selectivity across conditions because these differences might result from difference in how the encoding model was estimated (see Sprague et al., 2018 for further discussion of this issue).

### Cross-training analysis

For analyses in which we assessed whether the similarity of multivariate patterns representing one-item and two-item trials, we again partitioned each condition into three sets. We then trained the IEM on two sets of one item data and tested on a single set of the two item data.

## Statistical Analysis

### Modeling response error

Response error was measured as the number of degrees between the presented angular location and reported angular location, such that errors ranged from 0° (a perfect response) to ±180° (a maximally imprecise response). To quantify performance we fit a mixture model to the distribution of response errors for each participant using MemToolbox (Suchow et al., 2013). For one-item trials, we modeled the distribution of response errors as a two-component mixture model, comprising a von Mises distribution centered on the correct value (i.e., a response error of 0°), corresponding to trials in which the sample location was remembered, and a uniform distribution, corresponding to guesses in which the reported location was random with respect to the sample location. We obtained maximum likelihood estimates for two parameters: (1) the dispersion of the von Mises distribution (SD), which reflects response precision; and (2) the height of the uniform distribution (*P_g_*), which reflects the probability of guessing. For two-item trials we fit a three-component mixture model that also included an additional von Mises component centered on the location of the unprobed item, corresponding to trials in which participants mistakenly reported the location of the unprobed item (i.e. swaps, Bays et al., 2009). We obtained maximum likelihood estimates for the same parameters as in one-item trials, with one additional parameter (*P_s_*), which reflects the probability of swaps.

### CTF selectivity

To quantify the spatial selectivity of alpha-band CTFs, we used linear regression to estimate CTF slope. Specifically, we calculated the slope of the channel responses as a function of spatial channels after collapsing across channels that were equidistant from the channel tuned for the position of the stimulus (i.e., a channel offset of 0°). Higher CTF slope indicates greater spatial selectivity.

### Cluster-based permutation test

We used a cluster-based permutation test to identify when CTF selectivity was reliably above chance. This procedure corrects for multiple comparisons (Maris and Oostenveld, 2007; Cohen, 2014). To this end, we identified clusters in which CTF selectivity was greater than zero by a performing one-sample *t*-test (against zero) at each time point in the alpha-band analyses (or at each time-frequency point in the time × frequency analysis). We then identified clusters of contiguous points that exceeded a threshold of *t* = 1.703 (which corresponds to a onesided *p*-value of .05 for 27 degrees of freedom). For each cluster, we calculated a test statistic by summing all *t*-values in the cluster. We used a Monte Carlo randomization procedure to empirically approximate a null-distribution for this test statistic. Specifically, we repeated the IEM procedure 1000 times but randomized the positions labels within each block so that the labels were random with respect to the observed response at each electrode. For each iteration, we calculated CTF selectivity across time (or across time and frequencies), and identified clusters as described above. For each iteration, we calculated the test statistic for the largest cluster, resulting in a null distribution of 1000 cluster test-statistics. Finally, we identified clusters that had test statistics larger than the 95^th^ percentile of the null distribution. Thus, our cluster test was a one-tailed test with an alpha level of .05, corrected for multiple comparisons.

### Bootstrap resampling tests

We used a subject-level bootstrap resampling procedure (Efron and Tibshirani, 1993) to test for differences in CTF selectivity (measured as CTF slope) between the one-item and two-item conditions. We drew 100,000 bootstrap samples, each containing *N*-many observers sampled with replacement (where *N* is the sample size). For each bootstrap sample, we calculated the mean difference in CTF selectivity between conditions, yielding a distribution of 100.000 mean difference values. We tested whether the mean difference was significantly different from zero in either direction by calculating the proportion of values greater than or less than zero. We doubled the smaller value to obtain a two-tailed *p*-value. In cases where the mean difference for all bootstrap samples were to one side of zero, we report the *p*-value values as *p* < .00001. We deemed results to be reliably above chance if *p* < .05.

### Data and Code Availability

Data and code will be made publicly available at the time of publication.

## Acknowledgements

This work was supported by NIMH grant 2R01MH087214-06A1. We thank Janna Wennberg for assistance with manuscript preparation.

